# Deep learning enables direct HLA typing from immunopeptidomics data

**DOI:** 10.64898/2026.04.08.717021

**Authors:** Matteo Pilz, Jonas Scheid, Alina Bauer, Steffen Lemke, Timo Sachsenberg, Jens Bauer, Annika Nelde, Josua Stadelmeier, Axel Walter, Hans-Georg Rammensee, Sven Nahnsen, Oliver Kohlbacher, Juliane S. Walz

**Author notes:** These authors contributed equally to this work (Matteo Pilz, Jonas Scheid, Oliver Kohlbacher, Juliane S. Walz). Deceased author (Alina Bauer).

## Abstract

The immune system eliminates malignant and infected cells through T-cell-mediated recognition of peptides presented by human leukocyte antigen molecules. Mass spectrometry-based immunopeptidomics enables unbiased identification of naturally presented HLA-restricted peptides and has become central to the development of T-cell-based immunotherapies. However, immunopeptidomics data reflects the combined peptide presentation of multiple HLA alleles, and determining which allotypes are represented in this multi-allelic complexity remains an unmet computational challenge. Here, we introduce immunotype, a deep learning-based ensemble predictor for HLA class I allotyping directly from immunopeptidomics data. Immunotype integrates peptide and HLA protein sequence information through transformer encoders and a graph neural network, complemented by a curated mono-allelic reference of known peptide-HLA binding preferences. Immunotype achieves an overall accuracy of 87.2% at protein-level resolution across diverse tissues and thereby enables rapid, cost-effective HLA typing of large-scale immunopeptidomics datasets.

## Introduction

T-cell recognition of human leukocyte antigen (HLA)-presented peptides plays a key role in the immune surveillance of infectious and malignant diseases^1–3^. Various T-cell-based immunotherapeutic approaches aim to utilize respective peptides to therapeutically induce anti-tumor T-cell responses^4–6^. Mass spectrometry (MS)-based immunopeptidomics enables the direct identification of HLA-presented peptides and has significantly emerged with extensive investigation in preclinical as well as clinical studies^7–10^. However, peptide binding to HLA class I molecules is highly allele-specific^11^. Human individuals express up to six different classical HLA class I alleles, two each at the HLA-A, HLA-B, and HLA-C loci. HLA alleles are classified at different levels of resolution, with two-digit types distinguishing allele group and four-digit types distinct HLA protein sequences^12^. Each allele’s unique peptide-binding groove gives rise to allele-specific binding motifs, characterized by conserved anchor residues typically at the second position and the C-terminus of the peptide, resulting in donor-specific immunopeptidomes composed of distinct peptide subsets^11^. This multi-allelic complexity poses a major challenge in inferring HLA class I alleles from an individual’s immunopeptidome. Numerous computational approaches have been developed to resolve subsequent issues of this complexity by predicting peptide-HLA (pHLA) binding^13–15^ or deconvoluting allele-specific peptide subsets^16,17^ leveraging a combination of *in vitro* binding affinity data and immunopeptidomics-based eluted HLA peptides. These tools have been greatly enabled by mono-allelic cell line studies^18–20^ and public repositories such as the Immune Epitope Database^21^ (IEDB), the Peptides for Cancer Immunotherapy Database^22^ (PCI-DB), and the SysteMHC Atlas^23^, containing millions of peptides from thousands of benign and malignant samples thereby providing unprecedented insight into the complexity of antigen processing and presentation. While these efforts have improved pHLA binding predictors and enabled HLA allele-specific motif deconvolution, predictors modelling the full immunopeptidome complexity to infer the individual’s set of HLA class I alleles are lacking. Current HLA typing approaches, ranging from sequence-specific oligonucleotide probe hybridization and PCR amplification to high-resolution DNA and RNA sequencing^24–26^, remain labor-, time-, and cost-intensive, often leaving large immunopeptidomics datasets unannotated with HLA typing information. Here, we introduce immunotype, which is the first tool that provides accurate protein-level typing of HLA class I alleles directly from immunopeptidome data, enabling rapid, cost-efficient analysis of existing and newly generated datasets.

## Results

### Immunotype architecture and training strategy

Immunotype combines a graph neural network (GNN) including transformer layers with a lookup table from publicly available pHLA elution likelihood (EL) data designed to predict HLA class I alleles directly from immunopeptidomics data (Fig. 1a). To ensure immunotype learned from data reflecting immunopeptidomics experiments, the GNN model development was based on the PCI-DB+ training set that provides the complexity of immunopeptidomics datasets. While this immunopeptidomics resource provided the primary training material, we also leveraged binding affinity (BA) and mono-allelic elution likelihood (EL) datasets as sources of high-quality evidence for pHLA interactions^13,18–21^. These datasets were integrated into a staged pretraining strategy in which the model was first optimized on BA data, then on EL data, and finally on *in silico* generated immunopeptidomics samples alleles (Extended Data Fig. 1a). This progression gradually shifted the model from learning single peptide-single allele relationships toward handling sets of peptides and multiple HLA labels simultaneously to fully exploit the combinatorial structure present in the PCI-DB+ training phase. Sequentially pretraining the model led to gradually increased performance, specifically for HLA-C alleles (Extended Data Fig. 1b).

**Fig. 1:**
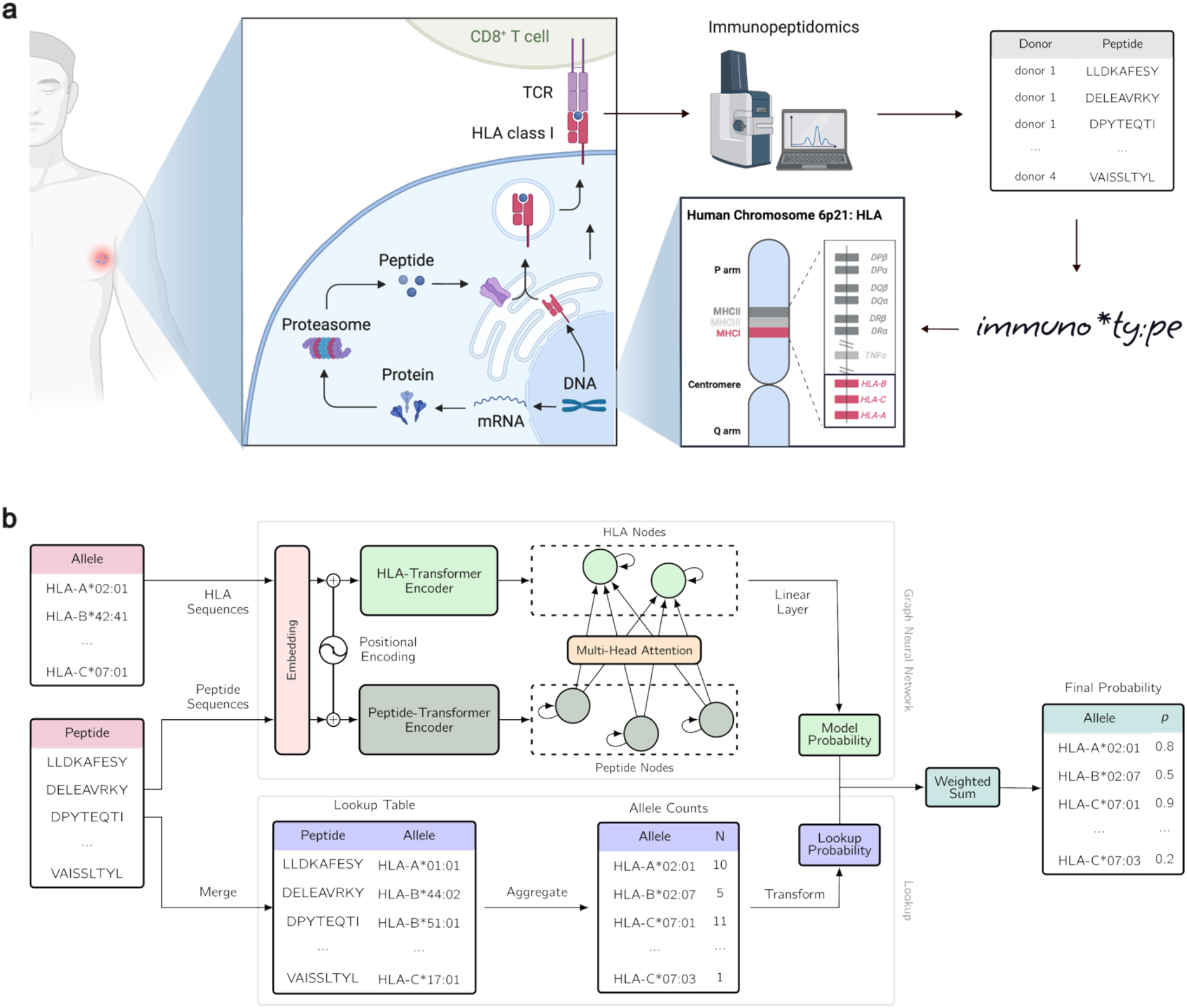
Immunopeptidomics and immunotype architecture overview. **a**, HLA class I-presented peptides arise from intracellular proteins degraded by the proteasome into short peptides, which are loaded on HLA molecules within the endoplasmatic reticulum. HLA-peptide complexes are transported to the cell surface and are recognized by the T-cell receptor of CD8^+^ T-cells, facilitating immune surveillance of infected and cancerous cells^*46*^. Mass spectrometry-based immunopeptidomics allows the direct identification of the landscape of HLA**-**presented peptides on the cell surface. Immunotype uses immunopeptidome data to infer HLA alleles of the respective donor. **b**, immunotype is based on a GNN and a lookup component and receives as input a list of peptides and alleles to infer the HLA alleles. The GNN learns HLA and peptide amino acid sequence representation separately with independent transformer encoder blocks after embedding them by the same layer. All peptide nodes in a sample are connected to all HLA nodes, and both are also connected to themselves. Two successive transformer graph convolutional blocks are applied for all nodes and use residual connections where the output of the first block is added to the output of the second. The final layer outputs a probability for each HLA allele. For the lookup component, peptide matches are searched, then the peptide count per allele is transformed and the top two scoring alleles per locus are obtained. The GNN and lookup operate independently to give a probability for each allele in the typing. These probabilities are weighted and added together to give the final output probability. Abbreviations: HLA - Human leukocyte antigen; CD - Cluster of differentiation; GNN - Graph Neural Network

The lookup component complements the GNN by leveraging allele-specific frequency patterns captured from a subset of high-confidence EL data to extract robust per-locus signals, particularly when peptide sets are small or highly skewed. As it relies on direct scoring from pHLA occurrences rather than learned contextual relationships, the lookup component provides a stable baseline.

To combine the strengths of both approaches, we integrated the GNN and lookup predictions into an ensemble method (Fig. 1b). Contributions were balanced through locus-specific weighting, ensuring that the ensemble favoured the component that performed best for each locus. The two best-scoring alleles per locus provide the predicted HLA typing labels.

### Evaluation and comparative HLA class I allele inference

Immunotype is a novel approach to predict HLA class I alleles directly from immunopeptidomics data. Several computational methods address aspects of immunopeptidomics data complexity, including pHLA binding predictors like NetMHCpan^13^ and motif deconvolution tools such as MHCMotifDecon^16^, but neither was designed for HLA typing. To allow a comparative evaluation, we implemented heuristic allele inference strategies (Extended Data Table 1) based on the tools binding prediction patterns and deconvoluted motifs, respectively.

We demonstrate that immunotype outperforms both the binding prediction and motif deconvolution tool heuristic at the level of individual HLA proteins with an accuracy of 87.2% over all HLA class I loci compared to 22.8% and 20.5%, respectively (Fig. 2a, Extended Data Table 2). At allele-group level resolution immunotype achieves an accuracy of 90.2%, exceeding 42.0% with binding prediction and 41.6% with motif deconvolution (Extended Data Fig. 3a). Immunotype’s prediction performance was also evaluated on an additional published immunopeptidomics dataset^27^ (n=17) confirming a mean accuracy of 91.2% for HLA-A, 94,1% for HLA-B, and 88.2% for HLA-C compared to the PCI-DB+ cross-validated accuracies of 92.5% for HLA-A, 87.4% for HLA-B, and 81.7% for HLA-C, respectively (Extended Data Fig. 3b, c).

**Fig. 2:**
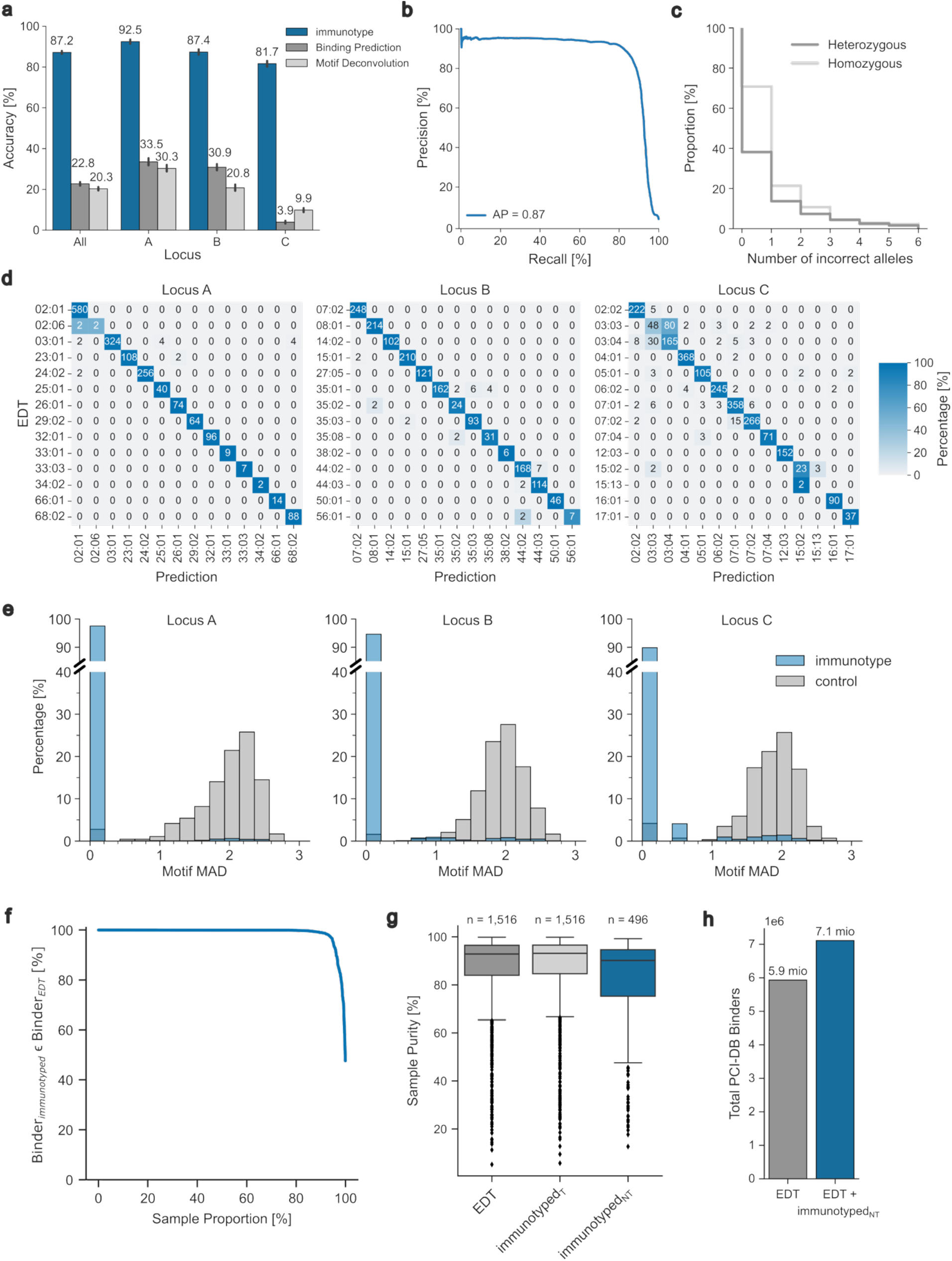
Immunotype performance on the PCI-DB+ dataset. **a**, Accuracy of immunotype, binding prediction, and motif deconvolution tool for locus A, B, and C on protein-level. **b**, Precision-recall curve of immunotype over all tested alleles and samples. **c**, Proportion of samples with number of incorrectly predicted alleles by immunotype for heterozygous and homozygous individuals. A donor is considered homozygous if one locus is homozygous. **d**, Most frequent unambiguous replacements with absolute numbers and percentage-based colouring between EDT and prediction. Predictions where both alleles are incorrect where ignored. **e**, Histogram of MAD between predicted alleles and EDT alleles for locus A, B, and C. The control was created by calculating the MAD over pairwise matches of all motifs. **f**, Per-sample predicted binders obtained with NetMHCpan and immunotype-defined (immunotyped_*T*_) alleles contained in predicted binders of EDT alleles, shown in relation to each sample’s peptide count. **g**, Per-sample ratio of predicted binders to total peptides (purity) for EDT alleles (n=1,516), immunotyped_*T*_ alleles, and non-EDT samples (n=496) typed with immunotype (immunotyped_*NT*_). **h**, Absolute binder increase of PCI-DB+ with EDT and immunotyped_*NT*_ samples. Abbreviations: Mean absolute motif deviation - MAD; EDT - Experimentally determined HLA typing

To further validate our strategy of selecting the two best-scoring alleles per locus for typing, we evaluated the precision-recall across all allele predictions, without any selection (Fig. 2b). Immunotype achieves an average precision of 0.87 across all predictions, indicating strong classifier performance even without restricting predictions to the top two alleles per locus. This demonstrates that the model preserves robust ranking capacity when allele selection thresholds are less stringent. Similarly, when analysing the top-k allele accuracy across all three predictors, the binding predictor and motif deconvolution tool improve substantially as *k* increases, whereas immunotype shows a high and stable trajectory that saturates early, underlining its reliable allele ranking capability (Extended Data Fig. 3d).

Due to the introduction of homozygosity labels, immunotype also predicts whether both parental copies carry the same allele at a given HLA locus. In our analysis, 55.2% (n=1,516) of samples were typed correctly (Fig. 2c), and in 1.7% of samples immunotype did not predict any correct HLA allele label. To infer whether a donor is homozygous or heterozygous at a given locus, we included additional labels indicating whether an allele could occur in single or duplicated form. While immunotype predicted homozygosity correctly in 74.8% of all cases for locus A, accuracy for locus B and C was markedly lower (B: 33.7%; C: 38.9%) (Extended Data Fig. 4a). To understand this discrepancy, we examined the allele probabilities obtained from the lookup component. We compared the distributions of the second-best lookup probability per allele at each locus between homozygous and heterozygous samples of the PCI-DB+ dataset (Extended Data Fig. 4b). For locus A, a clear separation between both distributions could be observed, however, loci B and C did not exhibit this behaviour.

We further investigated systematic patterns of prediction errors by quantifying the most frequently misclassified HLA alleles of immunotype (Fig. 2d). Misclassifications rarely occurred with alleles of locus A and B except for HLA-A*02:01, which was misclassified as HLA-A*02:06 in two of the four unambiguously identified cases. Errors were more frequent in HLA-C alleles and were largely confined to pairs of highly similar alleles. A prominent misclassification is HLA-C*03:03 and HLA-C*03:04, which differ by only a single amino acid outside of the binding cleft region and therefore does not affect the underlying binding motif. Misclassifications on allele-group level were rare, with one of the most frequent ones between HLA-B*55 and HLA-B*56 (Extended Data Fig. 5a). Systematic replacements, such as homozygous donors predicted as heterozygous or vice versa, were not observed (Extended Data Fig 5b, c). Next, we assessed the impact of all misclassifications by quantifying the mean absolute deviation (MAD) between binding motifs of replaced alleles and all pairs of alleles found in the PCI-DB+ to evaluate motif dissimilarity. In general, 97.7% HLA-A, 96.1% HLA-B, and 94.0% HLA-C misclassifications report a motif absolute difference < 1.0, indicating that most incorrectly predicted alleles are highly similar to experimentally-derived HLA typed (EDT) alleles (Fig. 2e).

To test the robustness of immunotype, we randomly selected peptide subsets in different sizes of each sample of the PCI-DB+ dataset and inferred the HLA typing from those peptides only. With only 20% of peptides of a sample immunotype reports an average accuracy of 84.3%. Performance drops drastically when the sample size was reduced to 1% with an average accuracy of 55.8% and an average of 42.5 peptides per sample (Extended Data Fig. 6a, b). We further evaluated immunotype performance under increasing typing complexity by introducing additional alleles into the prediction space that were not part of the training PCI-DB+ dataset. The prediction accuracy of 76.5% remains high with ten additional alleles and is gradually dropping with the introduction of additional alleles (Extended Data Fig. 6c)

Given the rise in large-scale immunopeptidomics datasets^28–30^ a rapid inference of HLA class I types is essential. We therefore quantified immunotype’s runtime under a range of common sample conditions. In a runtime benchmark on 100 samples, immunotype took on average 10.0 seconds per sample on CPU and 0.4 seconds on GPU (Extended Data Fig. 7a). Inference on samples including 20,000 peptides took about a minute on CPU and 6.5 seconds on GPU (Extended Data Fig. 7b). All runtimes were tested on a consumer-grade computer (see Methods), showcasing that immunotype is suitable for large-scale datasets without the need for dedicated high-performance hardware.

### Immunotype enables retrospective HLA typing and expands immunopeptidomics resources

We next investigated how HLA typing affects downstream immunopeptidomics analyses by comparing predicted binders from EDT alleles with those from immunotype-inferred alleles (immunotyped_T_). 75.3% of samples (n=1,142) contained the same predicted binders as in the EDT dataset due to the accurate prediction of HLA types by immunotype (Fig. 2f). Furthermore, losses of immunotyped_T_ binders compared to EDT binders could be largely attributed to a low sample input (Extended Data Fig. 8a).

Finally, we applied immunotype to annotate PCI-DB samples with missing EDT information (n=496, immunotyped_NT_) and compared their predicted binder to peptide ratios (sample purity) with both EDT and immunotyped_T_ alleles from the training dataset (n= 1,516, Fig. 2g). The purity of immunotyped_NT_ samples (median 90.4%) was comparable to EDT samples (median 92.9%) indicating that immunotype reliably infers HLA class I alleles and supports the inclusion of otherwise incomplete immunopeptidomics datasets. Inferring alleles of samples with missing EDT enabled the addition of 20,806 novel unique predicted binders to the PCI-DB, expanding the resource to a total of 448,499 unique HLA class I binders (Extended Data Fig. 8b). Overall, the absolute number of binders of the PCI-DB increased by 19.9% from 5,934,382 to 7,111,753 with the addition of 496 immunotyped_NT_ samples (Fig. 2h).

### Immunotype captures important properties of HLA sequences and HLA class I peptides

To evaluate whether the GNN component of immunotype captures biological properties of HLA sequences and their relationships with the HLA-presented peptides, we examined how the HLA sequence encoder represents HLA protein sequences in its latent space (Fig. 3a). The encoder shows a clear clustering of HLA loci and allele groups with similar HLA protein sequences. One of the most heterogeneous clusters contained alleles from six allele groups (HLA-B*14, HLA-B*15, HLA-B*38, HLA-B*39, HLA-B*67, HLA-B*73), where the majority shared common peptide binding motifs (Extended Data Fig. 9).

**Fig. 3:**
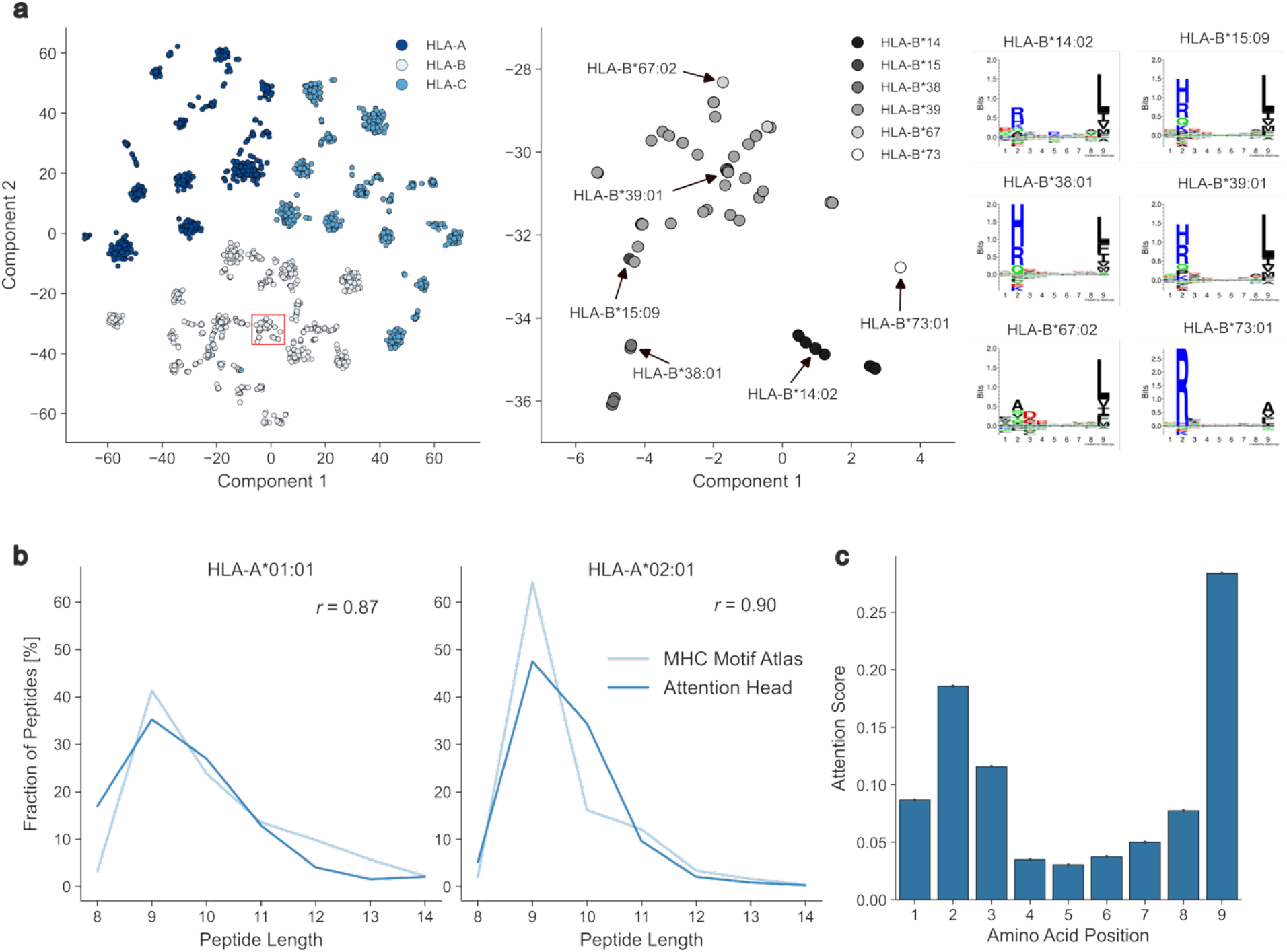
Immunopeptidome characteristics of immunotype’s neural network. **a**, t-SNE of MHC transformer encoder output displayed according to the HLA locus. A heterogeneous allele-group cluster (red square) is depicted in detail (right), highlighting the 6 most abundant alleles HLA-B*14:02, HLA-B*15:09, HLA-B*38:01, HLA-B*39:01, HLA-B*67:02, and HLA-B*73:01 with sequence logos created by Seq2Logo^*47*^. **b**, Correlation between peptide length preferences of MHC Motif Atlas and the MHC-peptide attention layer. Peptide length preferences from attention were correlated with individual heads with the strongest correlation from layer 2, head 3 (Extended Data Fig. 2). **c**, Attention scores returned by the peptide sequence transformer encoder for peptides with 9 amino acids. Abbreviations: MHC - Major histocompatibility complex

To investigate if immunotype learned well-known characteristics of HLA alleles and their associated binding motif properties, we compared the length distribution of HLA-A*01:01- and HLA-A*02:01-associated peptides (Fig. 3b). Using random peptide inputs, we extracted attention scores from HLA nodes given to each peptide and selected the ten peptides with the highest attention per allele. The resulting peptide length distribution correlates strongly with those from the MHC Motif Atlas^31^ (HLA-A*01:01 – *r=*0.87; HLA-A*02:01– *r*=0.90), similarly to the peptide correlation over all alleles in the training dataset (*r=*0.72, Extended Data Fig. 3), indicating that immunotype captures allele-dependent length properties of HLA class I alleles. This was further confirmed with the attention scores of the peptide sequence encoder showing a strong preference for position 2 and 9, the well-known binding anchor positions of 9mer peptides^11^ (Fig. 3c).

## Discussion

Technical and computational advancements in MS-based immunopeptidomics in recent years led to the rapid increase of large-scale datasets, enabling comprehensive antigen discovery for the development of T-cell-based immunotherapy approaches and contributing to the understanding of immunosurveillance mechanisms and autoimmune phenomena^7–10^. With the increasing availability of large-scale immunopeptidomics datasets, there is a growing need for rapid and accurate inference of HLA class I alleles directly from identified, HLA-presented peptides. Existing pHLA predictors of motif deconvolution tools, such as NetMHCpan^13^ and MHCMotifDecon^16^ can be repurposed to approximate HLA class I typing using predicted binding patterns or deconvolved motifs, however, these tools are not designed to resolve the full multi-allelic complexity of immunopeptidomics data. Immunotype is the first tool specifically designed to provide high-resolution HLA class I allotyping from immunopeptidomics data by learning both allele-specific protein sequence features and the combinatorial structure of multi-allelic peptide sets. Using an artificial neural network including transformer sequence encoding and graph attention layers together with an allele-frequency-based lookup, immunotype outperforms pHLA prediction and motif deconvolution tools. While NetMHCpan is an established tool for pHLA binding prediction, it struggles with multi-allelic HLA typing. This likely stems from representational constraints of NetMHCpan reducing MHC proteins to 34 binding-pocket residues, representing similar alleles as indistinguishable since they share the same amino acid sequence within their binding pocket. Our investigation of alternative aggregation strategies for both methods did not yield significant improvement, suggesting that these shortcomings might be to some degree rooted in sequence encoding, rather than prediction summarization. MHCMotifDecon and other tools, such as MHCflurry^14^, MixMHCpred^15^, or MoDec^17^, rely on similar architectural limitations and likely face the same issue. Consequently, immunotype’s ability to leverage full-sequence information is presumably critical for resolving high-resolution HLA types.

Due to the pretraining and training strategy, we showcased that immunotype tends to reflect known HLA sequence and HLA peptide properties via learned attention scores, such as binding motif similarities, peptide length preferences of specific HLA alleles, and prioritization of well-known anchor positions P2 and P9^11^. Nevertheless, attention weights can be noisy indicators and may not fully capture the complex biological relationships involved ^32,33^.

Through extensive prediction error analyses, we demonstrated that misclassifications of immunotype coincide predominantly with motif similarity between alleles. This is illustrated by the frequent misclassification of HLA-C*03:03 and HLA-C*03:04, whose identical binding-groove sequences result in indistinguishable peptide profiles^20^. Consistent with these observations, expanding the candidate allele pool reduces accuracy, as the inclusion of additional alleles with near-identical binding motifs increases the likelihood of indistinguishable peptide presentations. Importantly, these error sources have only a minimal impact on downstream immunopeptidomics analyses, enabling both retrospective HLA typing of datasets with missing EDT information and time- and cost-efficient HLA class I allotyping for large-scale immunopeptidomics studies.

Assessing immunotype’s ability to distinguish homozygous from heterozygous individuals showed clear differences between locus A and loci B and C. Homozygosity was identified with higher accuracy for locus A, while performance was reduced for loci B and C. Beyond differences in lookup counts, the underlying causes can only be speculated and may include higher motif similarity^19^, reduced peptide distinctiveness^20^, and lower HLA-C expression on the cell surface^34^ resulting in limited training data. An additional limitation of immunotype is a potential population bias in the PCI-DB+ training dataset, which could influence allele and peptide frequency estimates^35^. As most data originate from similar geographic populations, immunotype may inherit these sampling biases, potentially reducing performance in underrepresented populations.

Expanding the PCI-DB+ dataset with future studies is expected to reduce class confusion, particularly among highly similar alleles. Incorporating data from HLA class II alleles could further extend the scope of immunotype to enable CD4^+^ T-cell target definition, which is increasingly recognized as essential for orchestrating durable anti-tumor immunity^36^. Furthermore, high-resolution typing of non-classical HLA molecules like HLA-E offers significant potential for universal immunotherapy targets due to their minimal polymorphism and capacity to engage both T-cells and NK cells^37^.

Beyond HLA typing, the rich pHLA information contained in large immunopeptidomics datasets may also support additional predictive tasks. With modest architectural adaptations, immunotype could be extended to jointly predict binding affinity and elution likelihood for individual pHLA or pMHC pairs. Currently, these capabilities are lost due to sequential fine-tuning across datasets. Transitioning to a unified multi-task learning framework may preserve shared representations and enable simultaneous binding affinity, elution likelihood, and HLA typing within a single model.

Together, our results not only establish immunotype as the first high-performance allotype predictor but also underline its value as a reliable, robust, and cost-effective HLA typing tool to unlock the full potential of immunopeptidomics datasets, particularly in settings where EDT is unavailable.

## Material and Methods

### Human and non-human MHC protein sequences

HLA protein sequences were obtained from the IPD-IMGT/HLA database^38^ and non-human MHC sequences from the IPD-MHC^39^ database (IPD-IMGT/HLA: release 3.63.0 IPD-MHC: release 3.16.0.0, both accessed 26/02/04). For HLA, only confirmed alleles with a fully annotated coding sequence were used to achieve the highest possible distinction between them. The allele annotation was shortened to the HLA protein-level^12^, removing synonymous DNA substitutions and differences in non-coding regions without an effect on binding motifs. Remaining duplicate alleles were removed. To ensure consistent sequence representation, only proteins ranging from 200 to 450 amino acids in length were retained to remove proteins with potentially incorrect annotation and limit the amount of padding necessary, resulting in a final dataset of 10,394 unique MHC protein sequences.

### Dataset creation for pretraining of immunotype’s GNN component

Pretraining steps in the immunotype’s deep learning model component were conducted on three curated training datasets comprising binding affinity (BA), elution likelihood (EL), and *in silico* immunopeptidomics data, each annotated with HLA alleles (Extended Data Fig. 1). The BA training dataset from NetMHCpan^13^ (v4.1b) served as the first pretraining source. Entries were filtered to retain only those with a corresponding MHC protein sequence, resulting in 183,240 peptide-MHC binding affinity pairs.

The EL dataset was compiled from various immunopeptidomics studies that reported pHLA pairs including the EL dataset that was used to train NetMHCpan 4.1. HLA class I data from MHC Motif Atlas^31^ was added (accessed 26/02/06). Additionally, publicly available mono-allelic cell line datasets MSV000080527^18^, MSV000084172^19^, MSV000090323^40^, and PXD009531^20^ from MassIVE^41^ and PRIDE^42^ were reprocessed using MHCquant 2.6.0^28^. The search parameters for MHCquant were set according to the published settings for each study (Extended Data Table 3). Peptides identified below the peptide-level 1% false discovery rate threshold were incorporated into the EL dataset. Duplicate entries were removed, yielding the final EL dataset of 4,625,669 pHLA pairs.

The *in silico* immunopeptidomics dataset was built by systemically sampling pHLA pairs of the EL dataset to mimic a typical immunopeptidomics dataset. Specifically, all multi-allelic cell line datapoints were discarded, and up to 200 peptides per allele were combined, resulting in a single sample with protein-level class I typing containing a maximum of 1,200 peptides annotated with a maximum of six unique HLA class I alleles. This sample size was chosen because the model was trained on multiple samples in each batch to increase the learning stability in a limited GPU memory setting. While this number of peptides is significantly lower compared to an immunopeptidomics sample in the PCI-DB^22^ (median 3,049, Extended Data Fig. 10a), increasing sample size during training did not improve performance. To allow for the maximum number of different pHLA combinations, new batches were sampled randomly from the data in each epoch. Duplicated alleles per HLA locus were allowed to simulate homozygosity.

### Creation of the PCI-DB+ training dataset

To construct a comprehensive and state-of-the-art training dataset, we used the PCI-DB resource (*DB_release_240822_default*) with the addition of a recently published immunopeptidomics dataset (PXD076027) reanalysed by MHCquant 2.6.0 to ensure consistency in data processing. Only samples with complete sequencing-based protein-level HLA class I typing were retained. Peptide modifications and duplicate peptide entries within samples were removed, resulting in a total of 6,823,438 pHLA pairs and a sample median of 3,049 peptides (Extended Data Fig. 10a). Each donor is associated on average with 2.0 samples. Samples originating from the same donor shared on average 23.1% of peptides. The immunopeptidomics dataset contains 154 unique HLA class I alleles, 40 on locus A, 78 on B, and 36 on C (Extended Data Fig. 10b). In total, 747 donors were available with 103, 74, and 102 donors annotated as homozygous for HLA locus A, B, and C, respectively (Extended Data Fig. 10c).

### Architecture and training of immunotype components

Immunotype combines GNN-based with lookup-based HLA typing from peptide input data. The lookup-based approach identifies alleles per locus with the two highest lookup probabilities derived from a normalized and transformed lookup dataset. This dataset was constructed from a subset of the EL data by removing negative peptide-MHC pairs and entries originating from multi-allelic cell lines. Unique pHLA pairs were retained, resulting in 279,862 pairs in total. Lookup-based scoring was initially performed using

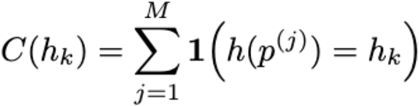

where *p*^*(j)*^ denotes the *j*-th peptide observed in the sample (*j = 1*, …, *M*), *M* is the total number of peptides in the sample, *h(•)* is a lookup function mapping a peptide to its uniquely associated HLA allele in the reference lookup table, *h*_*k*_ denotes the HLA allele under consideration, *1(•)* is the indicator function evaluating to 1 when the mapped allele equals *h*_*k*_ and to 0 otherwise, and *C(h*_*k*_) represents the total number of peptides in the sample assigned to allele *h*_*k*_ by the lookup procedure.

To enable the prediction of homozygous HLA loci, the same 6-fold cross-validation scheme on the donor-level as for the GNN was applied to the max-normalized, cubic-transformed lookup score of *C(h*_*k*_), yielding locus-specific thresholds for homozygosity. A cubic transformation was used to obtain a better separation between homo- and heterozygous HLA loci. The resulting thresholds *τ* for HLA-A, HLA-B, and HLA-C were *τ*_HLA−A_ = 0.425, *τ*_HLA−B_ = 0.3, and *τ*_HLA−C_ = 0.325, respectively. The final lookup-based probability *L(h*_*k*_) therefore was computed with

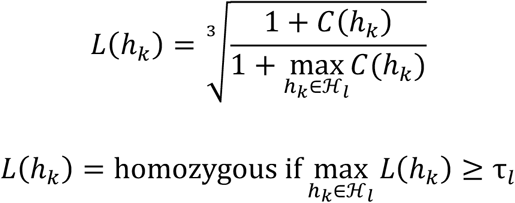

where ℋ_*l*_ denotes the set of all HLA alleles belonging to locus *l*.

The neural network in immunotype consists of two structurally distinct parts: the sequence encoder and the GNN, with a combined number of 5,053,057 parameters. The sequence encoders consist of a shared HLA sequence embedding with 128 dimensions, which was pretrained on non-human MHC as well. The encoding is done by two split transformer blocks with a feedforward dimension of 128/1,024, 8/8 heads, and 6/6 layers for the peptide/HLA-encoder. The GNN aggregates information over peptide sets using convolutional transformer layers with 32 output channels, 8 heads, and 2 layers with residual connections. In this architecture, each HLA allele is represented as an individual node initialized with its sequence-derived encoding, and each peptide forms its own node as well. Peptide nodes connect to themselves and to all HLA nodes, while HLA nodes also include self-loops. These pHLA connections allow the transformer layers to selectively focus attention on relevant peptides while down-weighting others. The GNN was implemented in Python (v3.10) with PyTorch^43^ (v2.0.0) and PyTorch Geometric^44^ (v2.3.0).

The GNN was trained and tested with a modified nested 6-fold cross-validation scheme on the level of tested donors on the PCI-DB+ dataset to obtain an unbiased estimate of generalization performance. The data were first partitioned into *k* outer folds. For each outer training fold, we conducted an additional inner split into training and validation sets (80/20%). The inner-validation scores were used exclusively to select hyperparameters. The model with the optimal hyperparameters was retrained on the entire outer-training fold and evaluated on the held-out outer test fold. The final performance is the average across the six *k* outer test folds. Hyperparameter tuning was done on the pretraining and fine-tuning datasets, if the architecture allowed it, but also validated and completed on the PCI-DB+ dataset. Fine-tuning steps were done sequentially (Extended Data Fig. 1a) after pretraining an autoencoder on the MHC and HLA protein sequences. This pretraining step was conducted to improve the model on alleles, which are not included in the remaining datasets. Together, those included a small proportion of the total number of non-human MHC and HLA proteins (0.6% MHC proteins, 6.4% HLA proteins annotated and confirmed). The model was fine-tuned on the BA dataset using the final model architecture, reusing only the MHC sequence encoder and embedding weights from the autoencoder stage. These BA and EL steps provided the model with detailed information on individual pHLA interactions.

The last fine-tuning stage mirrored immunotype’s intended application by supplying each HLA node with sets of multiple peptides from the i*n silico* immunopeptidomics data. We randomly generated 4,000 samples per epoch and continued training for 20 epochs until the performance plateaued. We applied sequential fine-tuning across stages to focus on the typing accuracy on the PCI-DB+ dataset computed on sample-level. All training steps were done with an Adam optimizer and a learning rate of 0.0001, on a machine with 16 VCPUs, 170 GB memory and two NVIDIA Tesla V100-SXM2-32GB.

### Immunotype ensemble predictor

The GNN and lookup predictions operate largely independent of each other. The input to both components is a set of peptides and the corresponding HLA alleles, which should be covered by the prediction. Both components provide a probability for each allele that was used to build the immunotype ensemble predictor. The same 6-fold cross validation was performed to obtain the optimal per-locus component probability weights. Probabilities were optimized with a weighted sum of one resulting in weights of 0.7/0.3 for HLA-A, 0.8/0.2 for HLA-B, and 0.8/0.2 for HLA-C (GNN/Lookup). Weights were tuned using the same nested cross-validation and grid search approach used for GNN hyperparameter optimization.

### Runtime analysis

The runtime of immunotype with default settings was tested on a consumer-grade computer running Microsoft Windows 11, with an NVIDIA RTX 4070 SUPER graphics card, an Intel i7-13700KF processor and 32 GB of memory. Single-sample runtime was measured for inputs containing 100-20,000 peptides, reflecting typical variability in immunopeptidomics experiments (Extended Data Fig. 10a). The prediction time of multi sample input ranging from 1 to 1,000 samples each containing 3,000 peptides with 8-14 amino acids was assessed on CPU and GPU.

### Peptide-HLA binding prediction

Peptide binding predictions for HLA class I was conducted using the nf-core/epitopeprediction pipeline^45^ (v2.3.1). NetMHCpan (v4.1b) was specified as the prediction tool. A length filter of 8-14 amino acids was set. HLA class I peptides were defined as binders with a percentile rank < 2.

### Benchmarking immunopeptidomics-based HLA typing

To benchmark immunotype we established baseline methods by adapting prediction scores of NetMHCpan (v4.1b) and MHCMotifDecon^16^ (v1.2b) to infer HLA typing. These scores were summed per allele, and the two alleles with the highest cumulative score per locus were taken as the predicted HLA types according to the following equation:

For a given HLA locus *l*, let *A*_*l*_ denote the set of candidate alleles. For each peptide *p* in the input set *P*, we identify the single allele *a* ∈ *A*_*l*_ that maximizes the tool-specific scoring function *S*(*p, a*), representing either the NetMHCpan *Score_BA* or MHCMotifDecon *RAW_SCORE*.

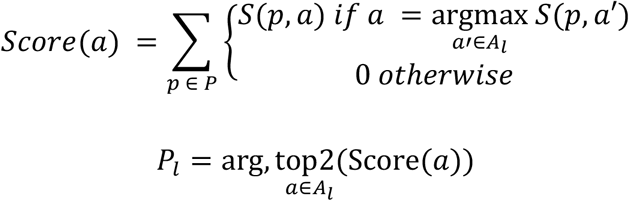

where *P*_*l*_ are the top2 alleles per locus *l* according to the *Score* function.

To evaluate the predictive performance of immunotype against NetMHCpan and MHCMotifDecon, a modified accuracy metric was utilized that accounts for the multiset nature of HLA typing. For each sample, let *L* denote the number of loci evaluated (*L=3* for HLA-A, -B, and -C). For a given locus *l*, let *T*_*l*_ represent the multiset of alleles from EDT and P_*l*_ represent the multiset of predicted alleles. The number of correct alleles at a single locus *m*_*l*_ was defined as the size of the intersection of these two multisets:

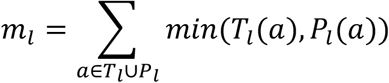

where *T*_*l*_*(a)* and *P*_*l*_*(a)* denote the count of allele *a* in the multiset *T*_*l*_ or *P*_*l*_. The overall accuracy is defined as the total number of correct predictions *m*_*l*_ across all loci normalized by the total number of EDT alleles *T*_*l*_.

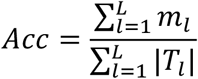

For a complete HLA class I typing across three loci, this metric yields seven possible discrete outcomes, ranging from zero to six correct alleles. As both NetMHCpan and MHCMotifDecon return multiple scores, we tested each score with different aggregation strategies (Extended Data Table 1).

## Data and Code Availability

Immunotype is available under the MIT licence as a web application on Hugging Face (https://huggingface.co/spaces/immunotype/immunotype), as a command-line tool on GitHub (https://github.com/AG-Walz/immunotype) and as a python package on the Python Package Index (https://pypi.org/project/immunotype). The training pipeline is available on GitHub (https://github.com/AG-Walz/immunotype-training-pipeline) alongside the pretraining MHC, BA, and EL and datasets. The PCI-DB+ data was compiled by complementing the PCI-DB data available at https://pci-db.org with the ProteomeExchange dataset PXD076027 (https://www.ebi.ac.uk/pride/archive/projects/PXD076027). Immunotype predictions of the PCI-DB are available in the new PCI-DB release *DB_release_260407_default*.

## Declarations

### Author’s contributions

M.P., J.S., O.K., and J.S.W. conceptualized this study. M.P., J.S., and S.L. curated the data in this study. M.P., J.S., and A.B. were involved in methodology and software development. T.S., J.B., A.N., H-G.R, O.K., and J.S.W. supervised this study. M.P., J.S, O.K., and J.S.W. wrote the manuscript. O.K., S.N., and J.S.W provided funding for this study. All authors read and approved the final manuscript.

## Acknowledgements

We thank M. Seybold from the Quantitative Biology Center (QBiC) for excellent technical support. We thank the de.NBI Cloud within the German Network for Bioinformatics Infrastructure (de.NBI) and ELIXIR-DE (Forschungszentrum Jülich and W-de.NBI-001, W-de.NBI-004, W-de.NBI-008, W-de.NBI-010, W-de.NBI-013, W-de.NBI-014, W-de.NBI-016, W-de.NBI-022), which provided computing resources to conduct the analyses. We acknowledge the use of BioRender.com to prepare certain figures. We acknowledge support from the Open Access Publication Fund of the University of Tübingen.

