## Extended Data Figures 1-10 for "Deep learning enables direct HLA typing from immunopeptidomics data"

### for

†Deceased author

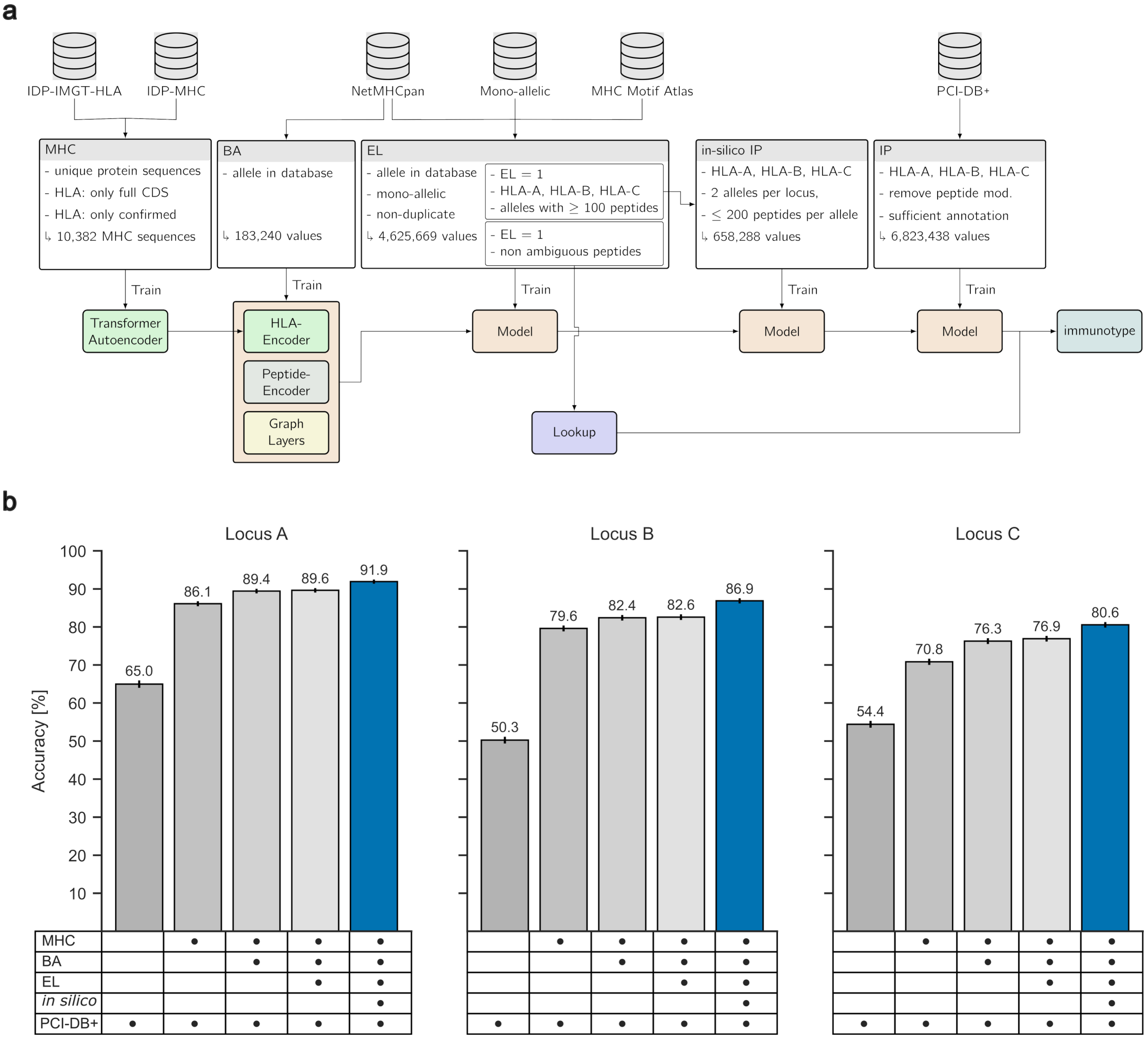


**Extended Data Fig. 1: Sequential training workflow and performance of immunotype**. **a**, Sequential training workflow of immunotype with corresponding data sources and properties. Pretraining of the immunotype neural network was initialized by training an autoencoder on 10,382 HLA and MHC protein sequences. Unconfirmed HLAs with incomplete coding sequences were discarded. The final model was subsequently trained on 183,240 BA values only including alleles with a corresponding protein sequence in the database. After filtering out for alleles in the database again and keeping only mono-allelic and non-duplicate entries, the model was afterwards pretrained on 4,625,669 EL values. One subset of the EL data serves as the lookup component, while another subset of 658,288 values was used to create the in-silico immunopeptidomics dataset. The final training step on the PCI-DB+ was done with 6,823,438 peptide-HLA values. **b**, HLA typing prediction accuracy on the PCI-DB+ dataset after different pretraining steps separated by locus. Abbreviations: HLA – Human leukocyte antigen; MHC - Major histocompatibility complex; BA – Binding affinity; EL – Elution Likelihood.

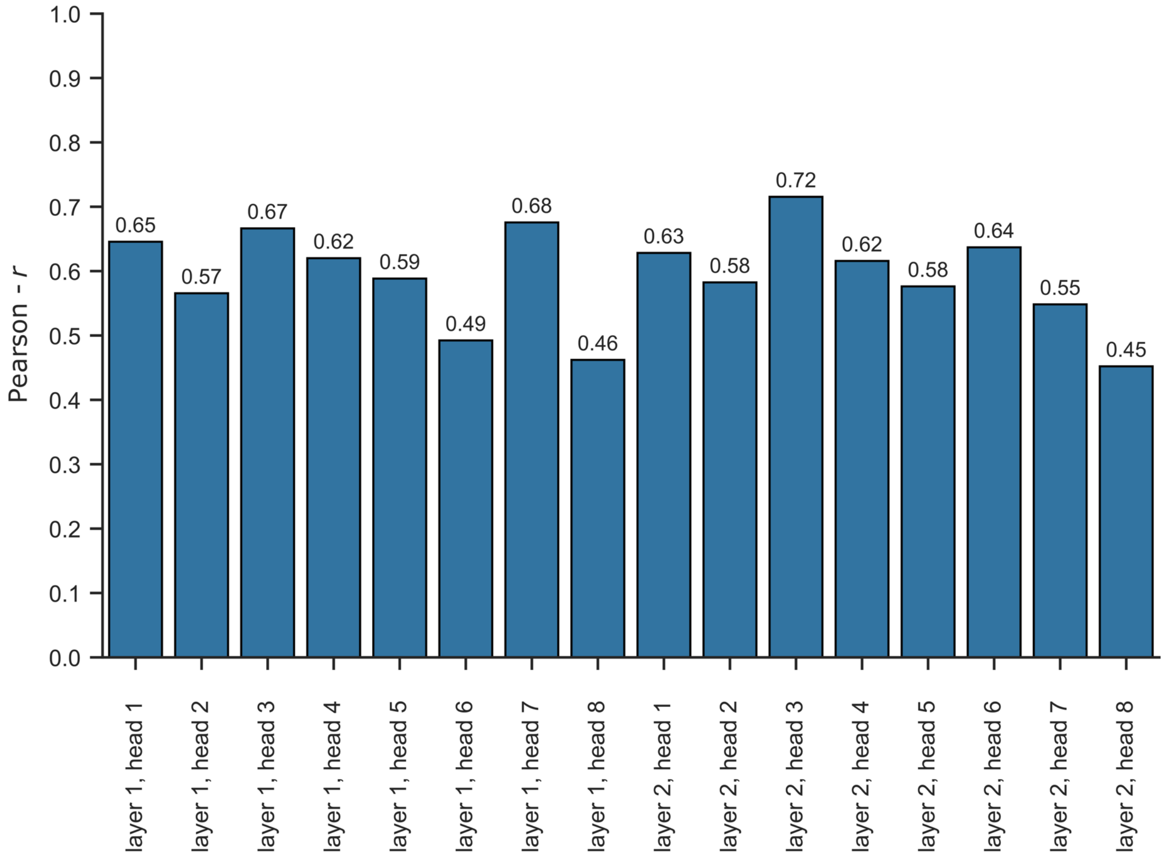


**Extended Data Fig. 2:** Pearson correlation coefficient between the peptide length preferences from all pHLA attention heads and published peptide length preferences from MHC Motif Atlas[31].

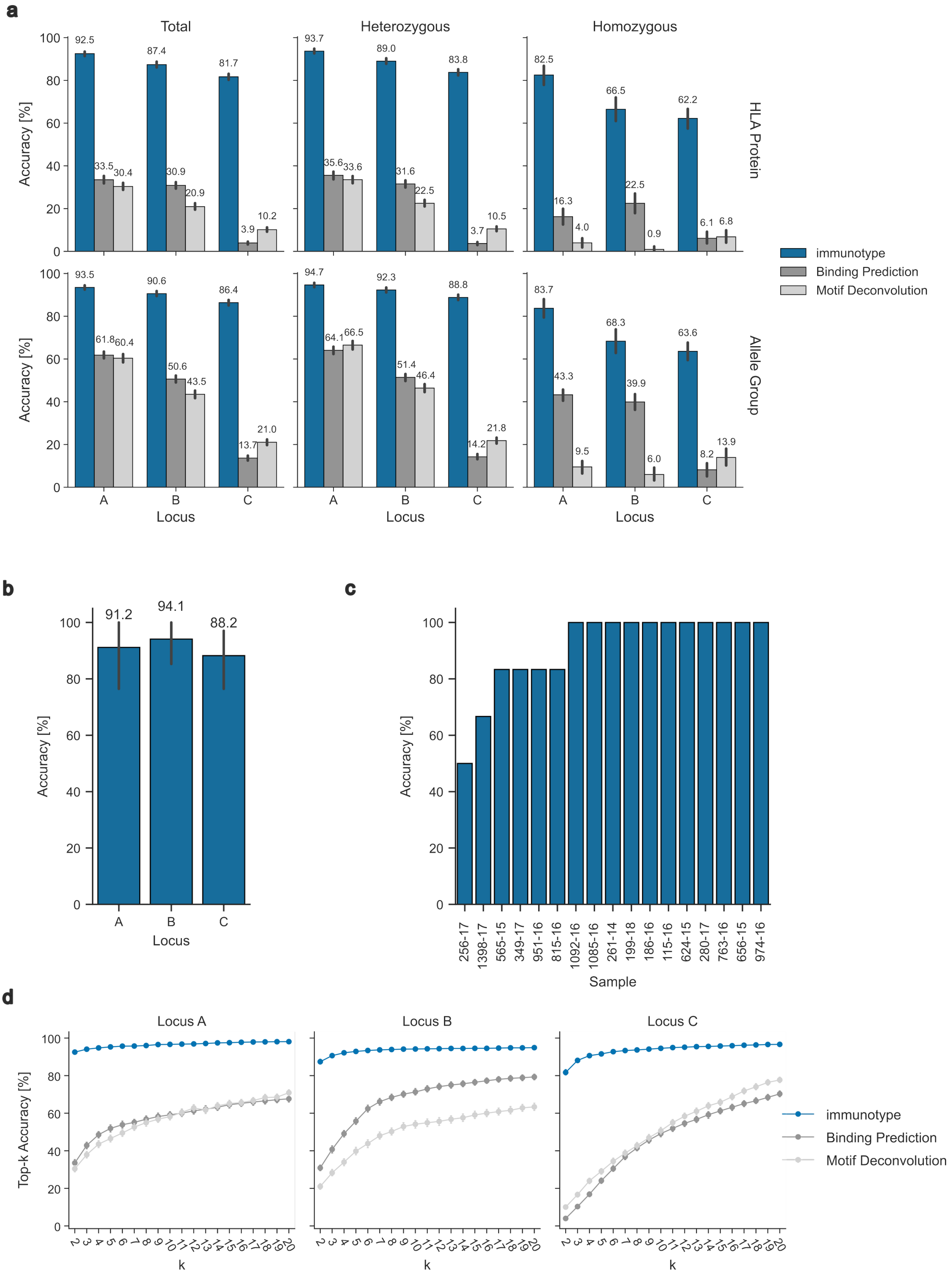


**Extended Data Fig. 3:** **a**, HLA typing prediction accuracy of immunotype, binding prediction, and motif deconvolution tools on the protein and allele-group level separated by locus. **b**, **c**, HLA typing accuracy of immunotype on public available dataset from Mühlenbruch et al.[27]

(n=17) per locus (b) and sample (c). **d**, Accuracy of HLA typings returned by the top-k alleles per locus from immunotype, binding prediction and motif deconvolution approaches.

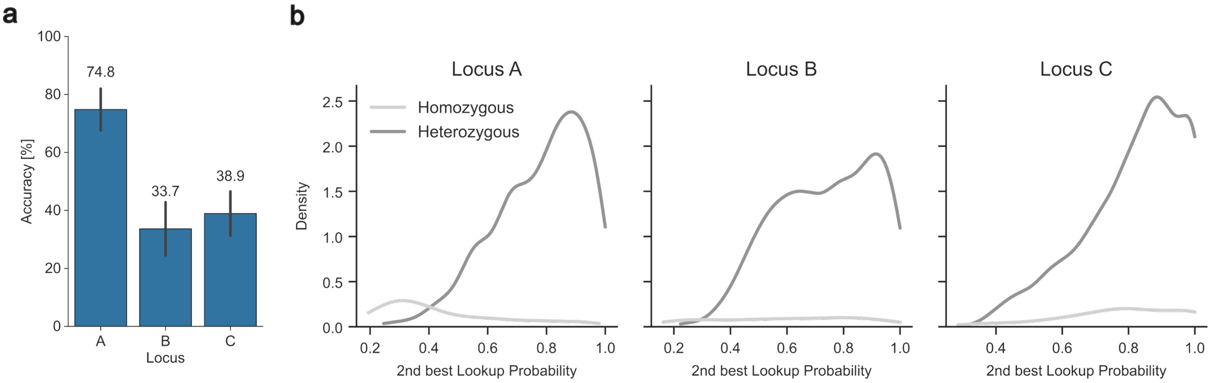


**Extended Data Fig. 4: Hetero- and homozygosity classification.** **a**, Accuracy of homozygosity prediction over locus A, B and C on PCI-DB+ dataset. **b**, Kernel density plot of alleles with the second-best lookup probability per locus.

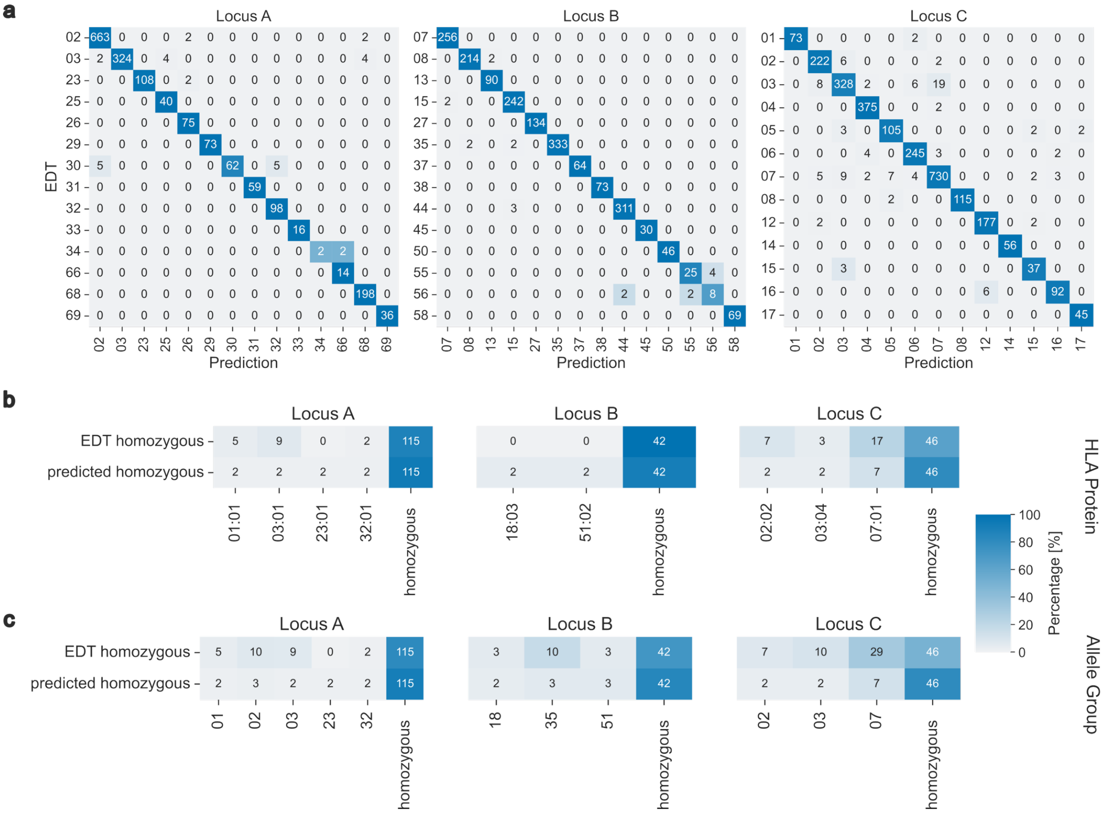


**Extended Data Fig. 5: Misclassifications analysis.** **a**, Frequency and absolute numbers of unambiguous replacements on allele-group level (**Error! Reference source not found.**d). **b**, **c**, Unambiguous replacements on the protein (c) and allele-group (d) level of samples from donors that are homozygous at the respective locus. The first row shows the replacement frequency and absolute numbers where the EDT was homozygous but was instead replaced by another typing. The second row shows the frequency and absolute numbers where immunotype predicted the allele to be homozygous but was instead another allele. Abbreviations: EDT - Experimentally determined HLA typing

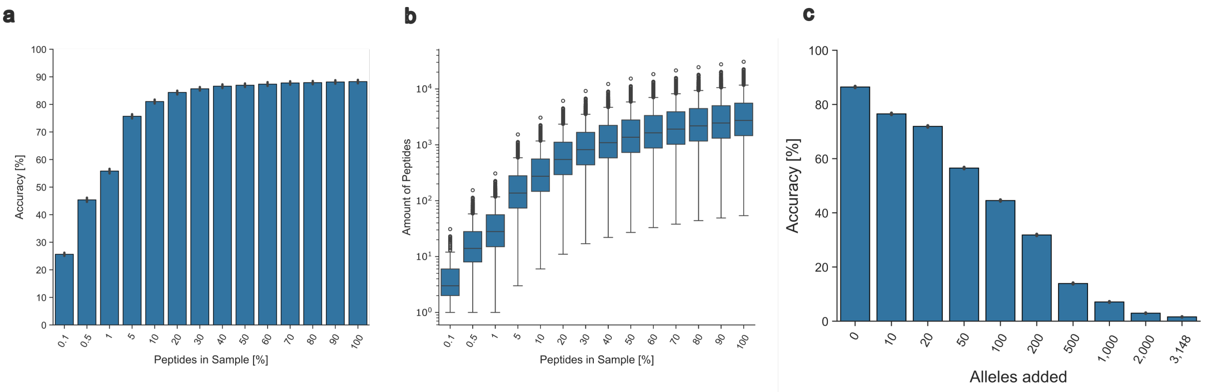


**Extended Data Fig. 6: Immunotype classification robustness. a**, Accuracy on the PCI-DB+ dataset after randomly selecting 0.1-100% of all peptides in a sample to predict the typing with immunotype with. **b**, Number of peptides in all samples after randomly selecting 0.1-100% of peptides in a sample. **c,** Bar plot depicting immunotype prediction accuracy with the random addition of alleles that were not part of the training dataset.

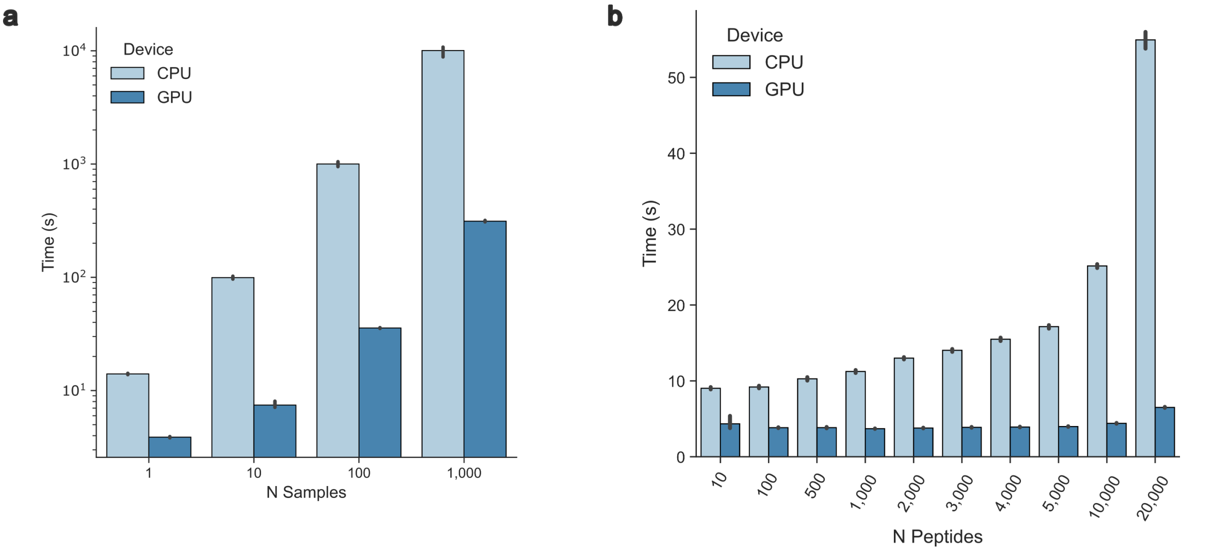


**Extended Data Fig. 7: Immunotype runtime analysis. a**, Prediction speed for N samples with 3,000 randomly generated peptides between 8-14 amino acids long on CPU and GPU. **b**, Prediction speed for a single sample with 10 to 20,000 peptides 8-14 amino acids long on CPU and GPU.

**a**

**b**

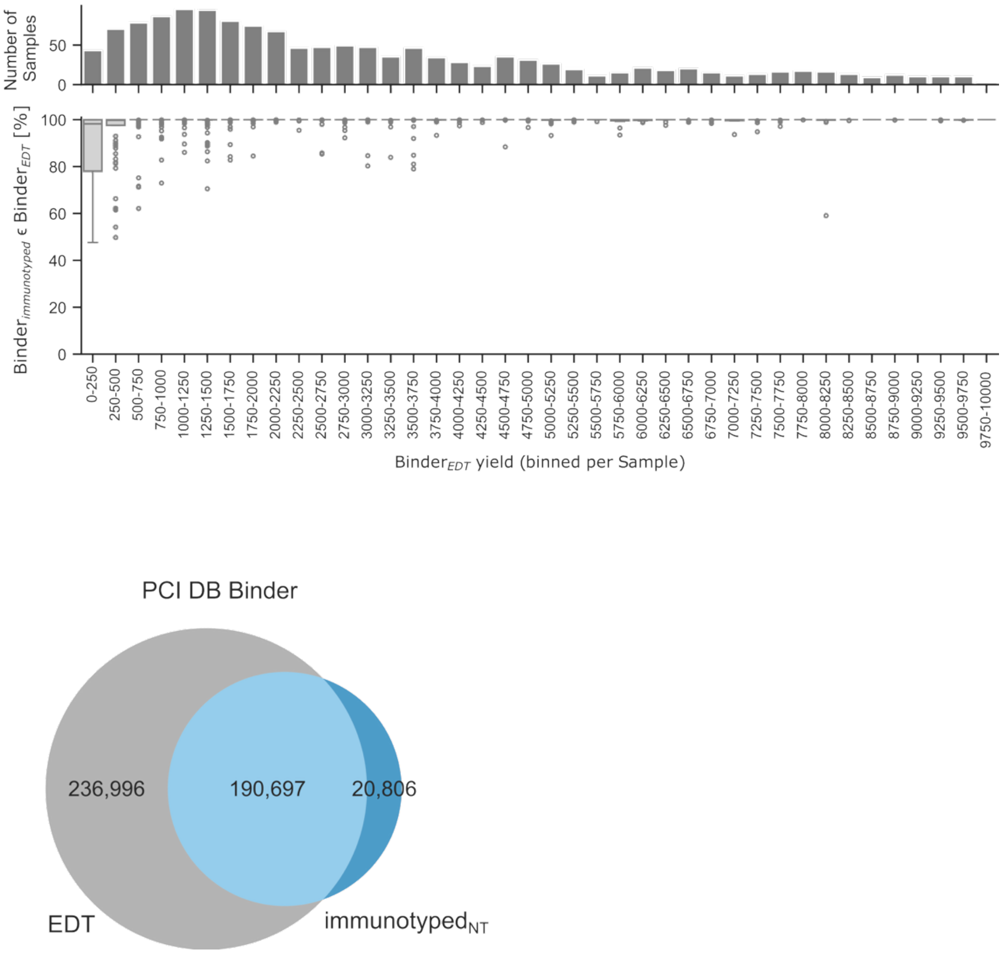


**Extended Data Fig. 8: a**, Per-sample predicted binders obtained with NetMHCpan and immunotype-defined (immunotyped$_{T}$) alleles contained in predicted binders of EDT alleles, shown in relation to the sample's binned binder yield (bottom) and the number of samples per bin (top). **b**, Venn diagram of NetMHCpan predicted HLA class I binders based on alleles from EDT and samples typed with immunotype where no EDT was available (immunotyped$_{NT}$).

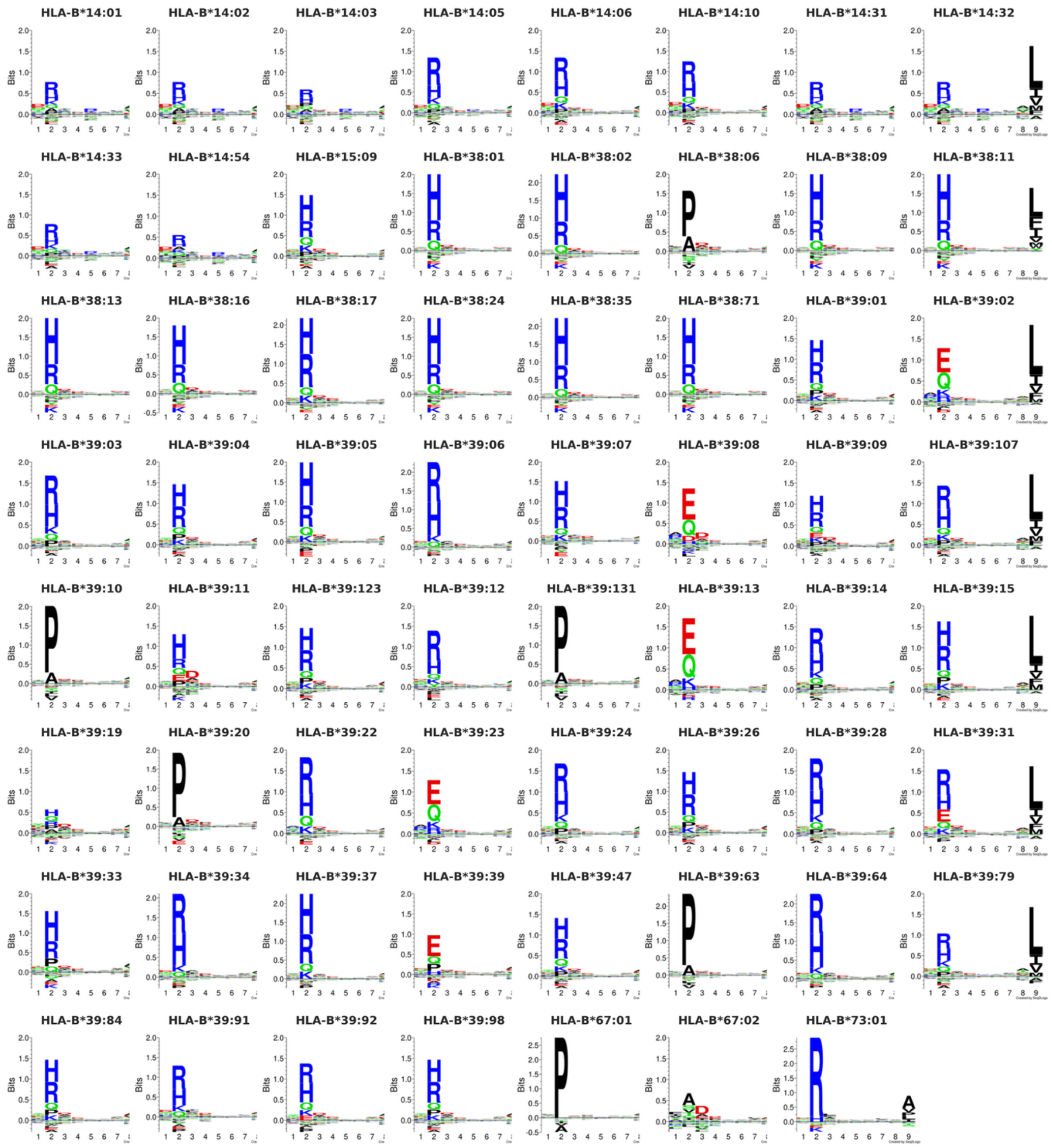


**Extended Data Fig. 9:** NetMHCpan available sequence logos of the heterogeneous cluster of Figure 3a generated by Seq2Logo[47].

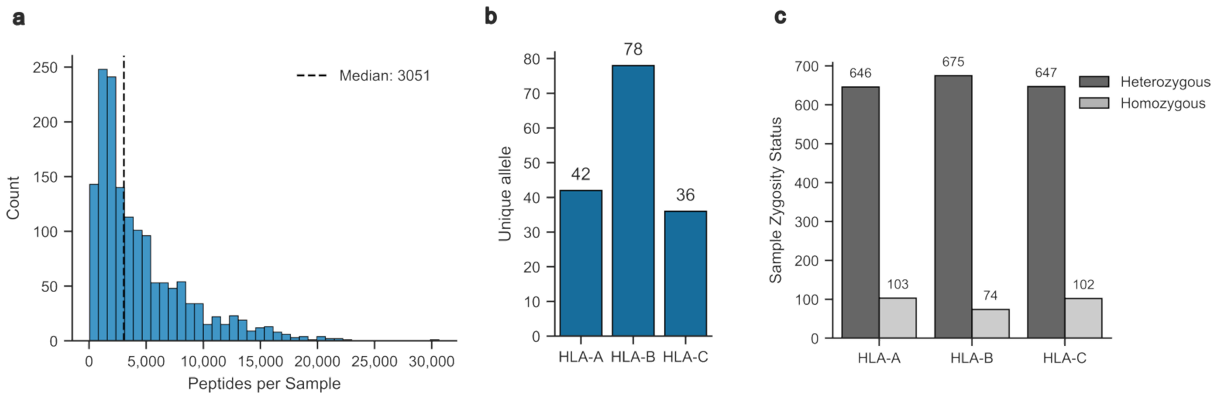


**Extended Data Fig. 10: Description of curated PCI-DB+ dataset. a,** Histogram displaying the number of peptides per sample. **b,** Number of unique alleles present per HLA locus. **c,** Number of samples with homozygous EDT per HLA locus. **d,** pHLA datapoints used during training shown per HLA allele and locus. Abbreviations: pHLA – Peptide-HLA complex.
